# TRAF3 Suppression Encourages B Cell Recruitment and Prolongs Survival of Microbiome-Intact Mice with Ovarian Cancer

**DOI:** 10.1101/2023.02.09.527814

**Authors:** Jonathan Zorea, Yair Motro, Roei D. Mazor, Yifat Koren Carmi, Ziv Shulman, Jamal Mahajna, Jacob Moran-Gilad, Moshe Elkabets

## Abstract

**Background:** Ovarian cancer (OC) has proven to be the most deadly of all gynecologic cancers. Clinical trials involving the use of immunotherapies in OC patients have produced disappointing outcomes, underscoring the necessity of identifying new immunomodulatory targets for the treatment of this cancer.

**Methods:** We conducted an *in-vivo* CRISPR screen of immunodeficient (NSG) and immune-intact wild type (WT) C57/BL6 mice to identify tumor-derived immune-escape mechanisms in a BRAC1- and TP53-deficient murine ID8 OC cell line (designated ITB1). To confirm gene expression and signaling pathway activation in ITB1 cells, we employed western blot, qPCR, immunofluorescent staining, and flow cytometry. Flow cytometry was also used to identify immune cell populations in the peritoneum of ITB1-bearing mice. To determine the presence of IgA-coated bacteria in the peritoneum of ITB1 -bearing mice and the ascites of OC patients, we employed 16S sequencing. Testing for differences was done by using Deseq2 test and two-way ANOVA test. Sequence variants (ASVs) were produced in Qiime2 and analyzed by microeco and phyloseq R packages.

**Results:** We identified tumor necrosis factor receptor-associated factor 3 (TRAF3) as a tumor-derived immune suppressive mediator in ITB1 cells. Knockout of TRAF3 (TRAF3KO) activated the type-I interferon pathway and increased MHC-I expression. TRAF3KO tumors exhibited a growth delay in WT mice vs. NSG mice, which was correlated with increased B cell infiltration and activation compared to ITB1 tumors. B cells were found to be involved in the progression of TRAF3KO tumors, and B-cell surface-bound and secreted IgA levels were significantly higher in the ascites of TRAF3KO tumors compared to ITB1. The presence of commensal microbiota was necessary for B-cell activation and for delaying the progression of TRAF3KO tumors in WT mice. Lastly, we observed unique profiles of IgA-coated bacteria in the ascites of OC-bearing mice or the ascites of OC patients.

**Conclusions:** TRAF3 is a tumor-derived immune-suppressive modulator that influences B-cell infiltration and activation, making it a potential target for enhancing anti-tumor B-cell responses in OC.

## Introduction

For many decades, ovarian cancer (OC) has consistently been ranked among the 10 most common and lethal cancers in women worldwide [1]. The majority of OC cases are classified as high-grade serous cancers, with approximately 50% of these cases involving alterations in homologous recombination (HR) repair genes [2]. Since most OC patients are diagnosed at stages III (51%) or IV (29%) [3], currently approved treatment modalities, such as surgery, chemotherapy, and/or targeted therapies (VEGF and PARP inhibitors, and the FRα antibodydrug conjugate), have contributed to an improvement in progression-free survival, but have not made a significant impact on overall survival [4]. The response rate to immunotherapies, including agents that block programmed cell death protein 1 (PD-1), its ligand PD-L1, or cytotoxic T-lymphocyte antigen 4 (CTLA-4), has also been low (5.9–12.3%) in OC patients [5,6]. Therefore, delineating new immune targets for the treatment of OC patients is urgently needed.

While clinical studies focusing on the role of T cells in OC have yielded disappointing results, the role of B cells in the OC tumor microenvironment (TME) has gained increasing attention [7]. The infiltration of regulatory interleukin (IL)-10-producing B cells into the ascites of OC patients is correlated with poor overall survival [8], while infiltration of immunoglobulin-producing plasma cells (IgGs [9,10] and IgAs [11]) has been positively correlated with improved patient survival. The importance of B cells in OC is thus gaining recognition, but it is still not fully understood how various factors affect the infiltration and activation of specific B cell populations in the TME.

It is known that B cells express IgA in response to antigen stimulation and costimulatory signals and that they function primarily against viruses and bacteria [12]. As such, bacteria–B cell interactions have been widely studied in various diseases (reviewed in [13]), including a study of breast cancer, in which bacteria coated with mucosal IgA were found to be significantly associated with estrogen levels [14]. In OC, a few studies have linked certain bacterial species to tumor progression in BRCA1-mutated OC patients [15,16], although the specific mechanism remains unclear.

In this study, using an ID8 murine OC cell line model, genetically modified with a loss of function in the *TP53* and *BRCA1* genes, designated here as ITB1, we identified a new tumor-derived immune suppressive mediator, tumor necrosis factor receptor-associated factor 3 (TRAF3). Through an unbiased targeted genetic screen, we showed that TRAF3 acts as an immunomodulator that promotes immune escape by suppressing the anti-tumor activity of B cells. We also showed that the anti-tumor activity of B cells is intimately associated with the gut microbiota.

## Materials and methods

### Cell line

A modified version of the ID8 mouse ovarian surface epithelial cell line, ID8-gTRP53-gBRCA1 (ITB1), was obtained from Dr. Iain A. McNeish [17] under an MTA agreement. This cell line was maintained at 37°C in a humidified atmosphere with 5% CO2 in DMEM medium supplemented with 10% fetal bovine serum (FBS), 1%L-glutamine, 200 mM, and 100 units each of penicillin and streptomycin. Mycoplasma infection was monitored regularly, and antibiotics (De-Plasma, TOKU-E, D022) were added to the culture medium as needed.

### gRNA pool library production

The murine ID8-gTRP53-gBRCA1-Cas9 cell line was generated by co-transfecting a lentiviral Cas9-Blast vector (Addgene #52962) with the packaging plasmids pCMV-dR8.91 and pCMV-VSV-G (Addgene #8454) into HEK293T cells. Following 48 h of transfection, the virus was harvested and its titer was determined. The Cas9-Blast lentivirus was then stored at −80°C until it was used for overnight infection of ID8-gTRP53-gBRCA1 cells. Selection with 5 μg/mL of blasticidin was performed two days later. To acquire clones with high Cas9 activity, single-cell sorting into 96-well plates was performed on the ID8-gTRP53-gBRCA1-Cas9 cells. Subsequently, infection with a lentivirus driving expression of a gRNA specific for CD274 and mCherry was carried out on multiple clones. Ten days post-infection, each clone was stimulated with 10 ng/mL of interferon (IFN)γ for 24 h, and the expression of PD-L1 was determined by FACS using an anti-CD274 antibody. The efficiency of Cas9 editing was determined by measuring the percentage of PD-L1 negative cells in the transduced population.

The mouse CRISPR Brie lentiviral pooled libraries consisting of 79,637 gRNAs were co-transfected with the packaging plasmids psPAX2 #12260 and pCMVVSV-G #8454 into HEK293T cells. Six hours after transfection, the medium was replaced with a medium supporting virus production (DMEM supplemented with 20% of FBS). After 48 h, the lentiviral medium was harvested and stored at −80°C. Before use in the *in-vivo* experiments, the ID8-gTRP53-gBRCA1-Cas9 cells were infected with the Brie lentivirus at a multiplicity of infection (MOI) of 0.06 and selected with puromycin for at least 10 days before injection, as described below in the section *‘In-vivo* experiments.’

### Generating knockout cell lines

To create knockout lines, the genome-scale CRISPR-Cas9 knockout (GeCKO) protocol was followed [19]. Oligos were designed using the same sequences as those provided in the gRNA library pool (described in ‘ gRNA pool library production’) and were phosphorylated and annealed with T4 PNK ligase. The lentiviral CRISPR plasmid was digested and dephosphorylated using the BsmBI restriction enzyme, and the annealed oligos were ligated into the lenti CRISPRv2 vector. The plasmid was transformed into stbl2 bacteria, and positive colonies were selected using ampicillin selection (100 μg/mL). The positive colonies were expanded, and plasmid DNA was isolated using the QIAGEN^®^ Plasmid Plus Midi Kit. The plasmid DNA was used to create lentiviruses and to infect ITB1 cell line to generate new lines expressing the desired genetic modifications. To this end, two TRAF3 guides, 5-CAGGTTCACGTGCTGTACCG-3 and 5-CGGTACAGCACGTGAACCTG-3, were used to created two stable ITB1 TRAF3 knockout (KO) clones, designated TRAF3KO1 and TRAF3KO2, respectively.

### Western blot analysis

All cells were harvested using a cell scraper and washed with ice-cold PBS. A lysis buffer supplemented with phosphatase inhibitor cocktails (BioTools, B15001A/B) and a protease inhibitor (Millipore Sigma, P2714-1BTL) was used to lyse the cells, which were then placed on ice for 30 min, followed by 3 min of ultrasonic cell disruption. The lysed cells were centrifuged at 14,000 rpm for 10 min at 4°C, and the supernatant was collected. The protein concentration was determined using the Bradford assay (Bio-Rad, Protein Assay, cat# 500-0006), and 1 μg/μL of total cellular protein from each sample was used to run a PAGE and transferred onto a PVDF membrane (Bio-Rad, 1704157). The membranes were soaked in a blocking solution of 5% BSA (AMRESCO, 0332-TAM) in TBS-T [Tris-buffered saline (TBS)-Tween 20 (0.1%)] for 1 h and then hybridized overnight with primary antibodies diluted in 5% BSA and 0.1% Tween. The following day, the membranes were incubated with horseradish peroxidase (HRP)-conjugated secondary antibodies (1:20,000, Jackson), diluted in the blocking solution. A chemiluminescent reaction was developed using enhanced chemiluminescence (ECL) (Westar Nova 2.0 Cyanagen XLS071.0250), and images were captured using an Azure Biosystems camera system. For nuclear and cytoplasmic separation, the Rockland Nuclear & Cytoplasmic Extract Protocol [18] was followed. In brief, the cells were lysed with a low-salt lysis buffer, which allowed the nuclear proteins to remain intact while the cytoplasmic proteins were lysed. Then, using a high-salt lysis buffer, the nuclear membrane and the nuclear proteins were lysed. The separation quality was verified by using an antibody against histone H3, which should be present only in the nuclear fraction.

### RNA isolation and real-time quantitative PCR

The isolation of total RNA and its conversion to cDNA were performed using the ISOLATE II RNA Mini Kit (Bioline, BIO-52073) and the qScript cDNA synthesis kit (Quanta Bioscience, 95047-100), respectively, as per the manufacturers’ protocols. A mix of cDNA and Fast SYBR qPCR Master mix (BioGate, EZ60) along with custom primer sets from IDT were used for qPCR analysis on a Roche light cycler 480 II machine.

### Flow cytometry

Isolated cells were blocked with anti-CD16/32 (anti-FcγIII/II receptor, clone 2.4G2) for 10 min at 4°C. Zombie Aqua™ fixable viability dye was then used to identify dead cells by incubating the cells with the dye for 15 min at room temperature. Surface markers were then stained by incubating the cells with the desired antibodies for 20–30 min at 4°C. The stained cells were analyzed using a CytoFLEX Flow Cytometer, and the results were analyzed using the CytExpert software.

For intracellular cytokine staining, cells were first cultured for 5–6 h with PMA (25 ng/mL), ionomycin (1 μM) and brefeldin A (5 μg/mL) to increase the intracellular cytokine staining signals. Surface markers were then stained, and the cells were fixed using 4% paraformaldehyde for 20 min at room temperature. The cells were permeabilized using a permeabilization buffer (1×) (eBioscience, 00-8333-56), and intracellular staining was performed by incubating the cells for 20 min. The cells were then analyzed on a CytoFLEX flow cytometer.

### *In-vivo* experiments

The *in-vivo* experiments were conducted using 6-to 8-week-old NSG mice (NOD.Cg-Prkdcscid Il2rgtm1Wjl/SzJ, Jackson labs) and wild-type (WT) C57/BL/6 mice (Envigo, Huntingdon, UK). The mice were housed in air-filtered laminar flow cabinets and provided with food and water ad libitum. Animal experiments were conducted in compliance with protocols established by the Institutional Animal Care and Use Committee (IACUC) of Ben-Gurion University of the Negev for ensuring animal welfare and minimizing discomfort. The animal ethical clearance protocol number used for the study was IL-23-05-2020(E).

Two different experiments were performed with the WT and NSG mice, as described in the Results section. In an experiment designed to investigate the role of anti-tumor immunity in the progression of ITB1-generated disease, WT and NSG mice were each injected intraperitoneally (i.p.) with 4-5 million ITB1 cells. In a separate experiment, which was designed to identify potential immunomodulators in ITB1 cells, WT and NSG mice were injected with 4-5 million ITB1 cells that expressed a gRNA library

Two experiments were performed only with WT mice. In the first, designed to examine the involvement of the microbiome in OC, one group of WT mice was pretreated with an antibiotic cocktail (ABX) for 2 weeks to deplete the total microbiome before TRAF3KO cell injection. This was achieved by adding ampicillin (1 g/mL), vancomycin (0.5 g/mL), neomycin sulfate (1 g/mL), and metronidazole (1 g/mL) to the drinking water and changing the water every two days [20]. The supplementation of the drinking water ABX was continued throughout the entire experiment. Two other groups of WT mice were injected with ITB1 or TRAF3KO cells and received normal drinking water (vehicle) throughout the entire experiment. For the FACS experiments, WT mice were sacrificed 2 weeks after the injection, and the peritoneal cavity was washed out with phosphate-buffered saline (PBS) to collect the contents.

For all the survival experiments, the body weight of the mice was monitored once a week, and the mice were sacrificed when a 20% increase in body weight was observed, indicating the development of ascites. The ascites fluid was collected for the isolation of bacteria or tumor cells. The digestive and reproductive systems were also collected for histology analysis.

### Bacterial DNA extraction

Bacterial DNA was isolated from ascites fluid or PBS wash using a modified phenol-chloroform protocol [21]. In brief, samples were lysed with a lysis buffer containing phenol, chloroform, and isoamyl alcohol (IAA) and then homogenized by bead-beating with bacterial disruption beads (RPI, 9833) for two cycles of 1 min each. The aqueous and phenol phases were separated by centrifugation, and the aqueous phase was re-suspended in IAA in a sterile tube. The bacterial DNA was then concentrated and isolated using isopropanol precipitation and ethanol washes. To minimize contamination, sterile conditions were used, and negative controls were taken by sampling the environment and collecting tools.

### Sorting of IgA+ and IgA-bacteria

The above-described PBS washes were collected from co-housed mice and then centrifuged at 8,000 ×g for 5 min at 4°C. The pellet was then resuspended in 100 μL of a blocking buffer containing 20% normal rat serum and incubated for 20 min on ice. Thereafter, the samples were stained with 100 μL of staining buffer containing phycoerythrin (PE)-conjugated anti-mouse IgA (1:12.5; eBioscience clone mA-6E1) for 30 min on ice. Following this step, the samples were washed 3 times with 1 mL of staining buffer. The anti-IgA-stained bacteria were then incubated in 1 mL of staining buffer containing 50 μL of anti-PE magnetic activated cell sorting (MACS^®^) beads (Miltenyi Biotec) for 15 min at 4°C. Finally, the samples were washed twice with 1 mL of staining buffer at 8,000 ×g for 5 min at 4°C, and sorted using MACS. The IgA-positive and the IgA-negative samples were then stored at −80°C for future use.

### Sequencing and sequence analysis

The genome-scale CRISPR-Cas9 library was sequenced on an Illumina MiSeq platform using a 250-bp paired-end chip. The sequences were mapped using a modified version of the caRpools R-package [22]. Differentially expressed guides were identified by loading the normalized gRNA count table into Deseq2 and determining the top differential genes based on the mean log2 fold change and false discovery rate (FDR). To identify significant pathways that were enriched or depleted in the screen, gene set enrichment analysis, including genes with a threshold of log2 fold change <(−1), was conducted using the ReactomePA [23] R package.

To amplify the V3-V4 region of the bacterial DNA isolated from the ascites of mice, a 2-step PCR-based protocol published by Holm et al. [24] was followed. The amplicons were then sequenced using the Illumina MiSeq platform (MiSeq Reagent Kit v2, 250-bp paired-end). Quality-checking of the demultiplexed sequencing reads was carried out using FastQC (Version 0.11.8) [25] and MultiQC (Version v1.7) [26]. Adapters, primers, and low-quality reads were removed using Fastp (Version 0.2, defaults parameters) [27]. The sequences were denoised by using Qiime2 version 2021.11 [28], and an amplicon sequence variant (ASV) table was created with taxonomic assignment using the SILVA database version 138, 99% NR [29]. Contaminant sequences were identified and removed based on control samples and sequence prevalence using the decontam package (Version 1.12) [30] and squeegee tool (Version 0.1.3) [31]. Further analysis of the ASV table was done using the MicroEco package (Version 0.11) [32] and phyloseq package (Version 1.38) [33] in R (Version 4.1.1).

A similar proof-of-concept analysis was performed for ascites from women with OC (see ‘Results section’).

### Statistical analysis

The *in-vitro* experiments were repeated a minimum of two to three times, with representative data/images being presented in the ‘Results’ section. For the *in-vivo* experiments, a minimum of 5 mice per group were used. Statistical analysis was conducted using GraphPad Prism 9 software and results are presented as means ± SEM. For experiments with two or more groups, a two-way ANOVA with Tukey’s multiple comparison tests was used. Significance levels of p-values of 0.05, 0.01, 0.001, or 0.0001 were calculated and are indicated in the figures by *, **, ***, or ****, respectively.

## Results

### CRISPR screen identifies TRAF3 as an immunomodulator of BRCA1-mutated OC

To investigate the role of anti-tumor immunity in the progression of ITB1-generated disease *in vivo*, we injected ITB1 cells i.p. into both immunocompromised NSG and immunocompetent WT mice and then monitored the survival of these tumor-bearing mice (Fig. 1A). The ITB1 cells did not form a single tumor mass, but rather caused a spreading disease in the peritoneum (as can be seen by immunohistochemistry staining), which also generated a significant accumulation of ascites (Fig. S1A). Although there was a significant difference in the rates at which the WT and NSG mice developed malignant disease, the median survival differed by only 8 days (41 vs. 49 days median survival). We, therefore, concluded that the ITB1 cells may have intrinsic immune escape mechanisms that suppress anti-tumor activity in WT mice, enabling rapid tumor progression.

**Fig. 1.**
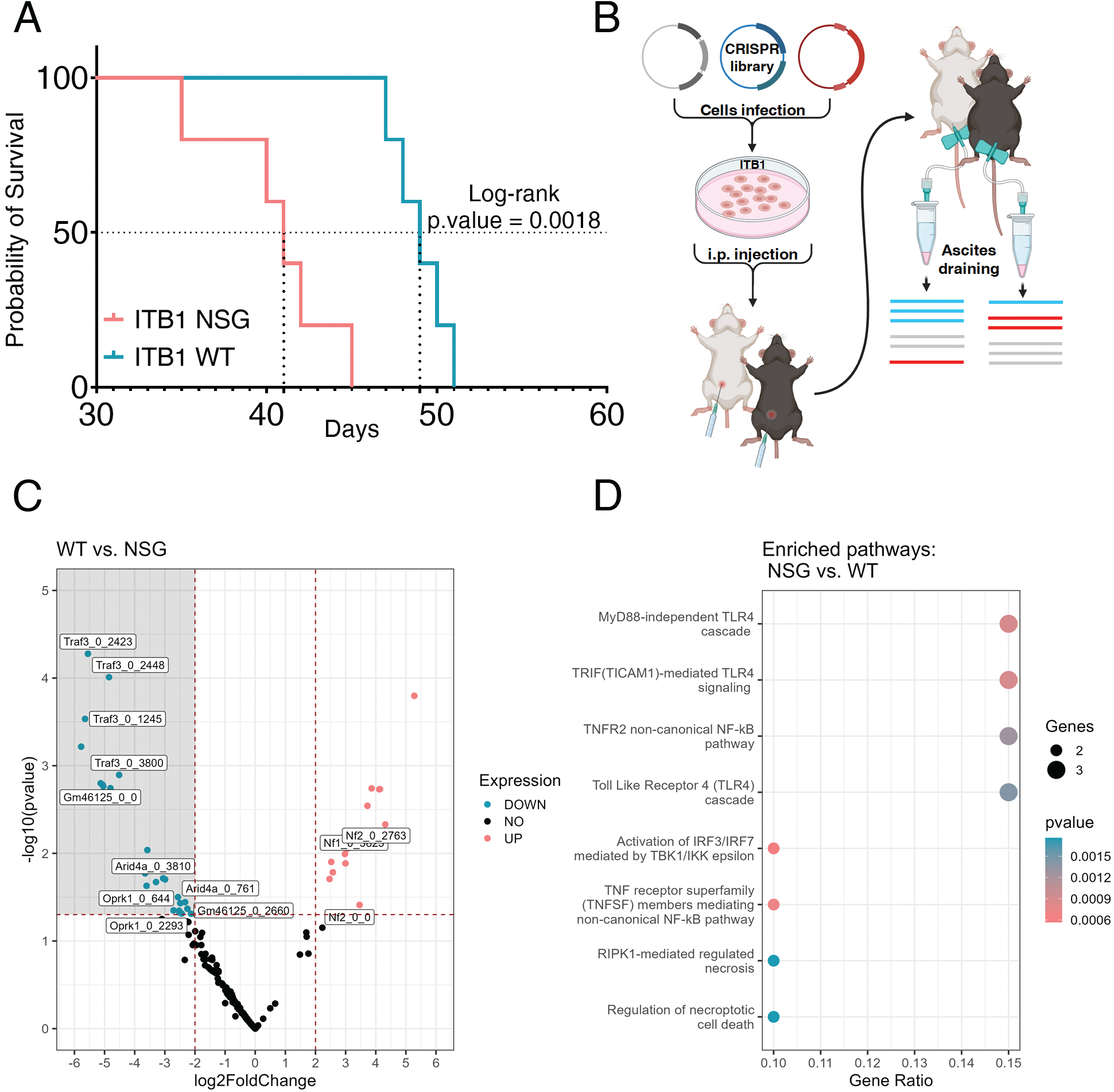
Identification of TRAF3 as a potential immunomodulator. **A** Kaplan-Meier plot showing the survival of two groups of ITB1-injected mice: pink line - NSG mice and cyan line - WT mice. **B** Schematic representation of library preparation, injection, and library DNA isolation. **C** Volcano plot showing enriched guides in ITB1-injected WT mice (pink) compared to ITB1-injected NSG mice (cyan). Guides that are significant appear above the horizontal dotted line. **D** Reactome pathways enriched with the enriched guides in the NSG group. The size of the dots represents the number of genes related to the pathway, and the color represents the p-value.

To identify potential immunomodulators in ITB1 cells that may be responsible for such immune escape, we knocked out a panel of immune-related genes by using a targeted guide RNA library of known immunomodulators [34] and then performed an *in-vivo* screen in WT and NSG mice (Fig. 1B). Specifically, we sought to identify genes that would facilitate the elimination of the tumor cells in the immunocompetent WT mice but not in the NSG mice. To this end, we first transduced the ITB1 cells with a lentiviral vector that expressed Cas9 and generated single-cell clones with the aim of identifying clones with a Cas9 editing efficiency of at least 95%. To test the editing efficiency, single clones overexpressing gRNA targeting PDL1 following IFNγ stimulation were stained for PDL1 (Fig. S1B). Clone 8, identified with an efficiency of 96.5%, was expanded and transduced with the complete gRNA library. The distribution of the library was verified by sequencing the amplicons produced from this ITB1 library-expressing clone. Out of 3878 guides, consisting of 1878 guides targeting 313 immune-related genes (six guides per gene) and 2000 guides as a control, only 500 guides were sequenced less than four times. After confirming that the majority of the guides were sequenced more than 10 times (Fig. S1C), we injected the gRNA library transduced ITB1 cells i.p. into NSG and WT mice. As soon as the production of ascites had resulted in a 20% increase in the weight of the mice, we drained the ascites, collected the cells, and isolated the total DNA for gRNA sequencing and analysis (Fig. 1B). By comparing the expression of gRNAs in the tumor cells growing in NSG and WT mice, we found six significantly differentially expressed genes in NSG mice compared to WT mice. Of these, two genes, NF1 and NF2, were significantly increased in the WT mice, and four genes, ARID4A, GM46125, OPRK1, and TRAF3, were significantly increased in the NSG mice (Fig. 1C). Notably, we included only those genes that were represented by two or more guides (out of the six represented in the library).

In view of our search for new tumor-derived targets that could sensitize ITB1 cells to anti-tumor immunity in mice, we focused on gRNAs and their signaling pathways that were increased in NSG mice compared to WT mice. We therefore conducted a Reactome pathway analysis on all genes whose absolute log2 fold change value was higher than one. The analysis revealed that TRAF3 was involved in six of the eight enriched pathways shown in Fig. 1D. Since TRAF3 had the highest expression difference and has been identified as a negative regulator of major histocompatibility complex class I (MHC-I) proteins and as being associated with a poor response to immune checkpoint blockade [35], we decided to further investigate its role in mediating immune escape in the ITB1-OC model.

### TRAF3 depletion activates the type-I interferon pathway and MHC-I expression and enhances the immunogenicity of ITB1 tumors

To determine the effect of tumor-derived TRAF3 on immune escape, we created two stable ITB1 TRAF3 knockout (KO) lines, designated TRAF3KO1 and TRAF3KO2 (Fig. 2A). We first compared the proliferation and migration abilities of ITB1 and TRAF3KO1/2 cells *in vitro* and observed a slight reduction in the proliferation rate of TRAF3KO1/2 cells, but no differences in their ability to close the scratch in a wound-healing assay (Fig. 2B and C). As TRAF3 is known to regulate the NF-κB pathway [36], we compared the nuclear protein levels of p65 and RelB, the total levels of p100 and p52, and the mRNA levels of IkBa, IL6, and RelB in ITB1 and TRAF3KO1/2 cells. We observed no significant activation of the NF-κB pathway in TRAF3KO1/2 cells, except for a minimal activation of the non-canonical NF-κB pathway, as indicated by Relb expression (Fig. S2A). Since TRAF3 is also known to mediate type I IFN signaling [36], we compared the mRNA levels of several IFN-stimulated genes in ITB1 and TRAF3KO1/2 cell lines and found a significant increase in the two TRAF3KO cell lines in the levels of three major genes in the type-I IFN pathway, namely, IFN-β, ISG15, and IRF7 (Fig. 2D). We also found the upregulation of type-I IFN to be associated with activation of the stimulator of interferon genes (STING) pathway, as indicated by the nuclear levels of TBK1 and STING upon western blot analysis (Fig. 2E and Fig. S2C) and the nuclear levels of STING upon immunofluorescence analysis (Fig. 2F and Fig. S2D). The strong activation of type-I IFN in TRAF3KO1/2 cell lines and the known association of TRAF3 with MHC-I expression and the activation of tumor immunity prompted us to examine the levels of MHC-I expression on the surface of the tumor cells. Indeed, TRAF3KO1/2 cells expressed higher levels of MHC-I compared to ITB1 cells, and the expression of MHC-I on the surface of TRAF3KO1/2 cells was almost at the same as that of IFN-stimulated ITB1 cells (Fig. 2G).

**Fig. 2.**
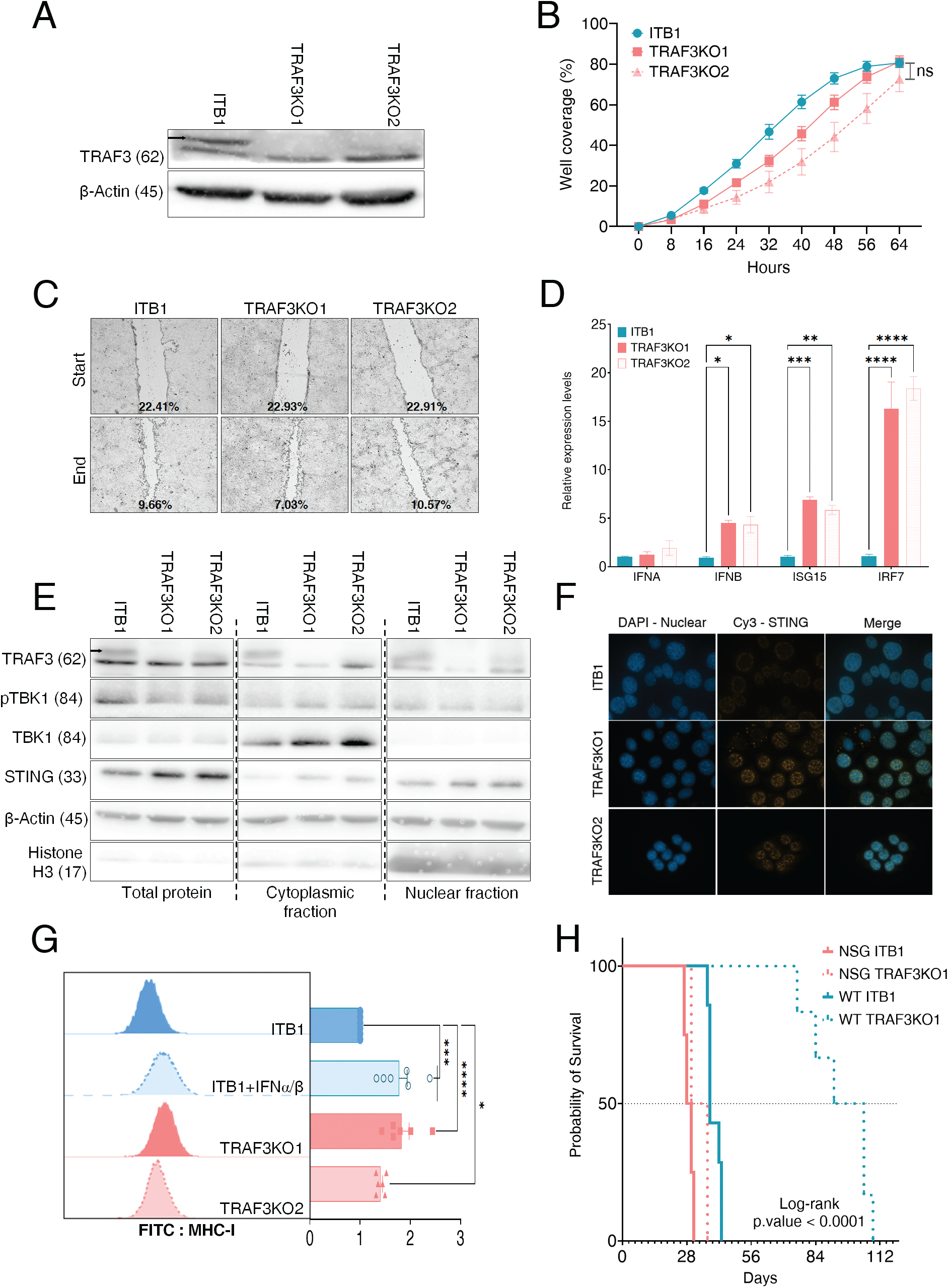
TRAF3 knockout (TRAF3KO) cells activate type-I interferon (IFN)-related pathways. **A** Western blot of TRAF3KO clones demonstrating the depletion of TRAF3 compared to the wild-type (WT) TRAF3 of ITB1 lysates. **B** Growth of ITB1 and TRAF3KO cells in culture was monitored every 8 h until confluency. The cyan line represents the growth of ITB1 cells, and the pink lines represent the growth of TRAF3KO cells. **C** Scratch assay showing the migration of all cells over 48 h into the scratched area. Data was acquired using a live cell imaging system (Julistage). **D** Quantitative PCR shows the mRNA levels of genes related to the type-I IFN pathway (IFNA, IFNB, IRF7, IFIT1, ISG15). Cyan and pink bars represent the levels in ITB1 and TRAF3KO cells, respectively. **E** Western blot of proteins in the STING pathway in ITB1 and TRAF3KO lysates. Actin and histone H3 were used as loading controls. Numbers in parentheses represent the protein size in kDa. **F** Immunofluorescence images showing the expression of STING (Cy3) in the nucleus (DAPI) of ITB1 and TRAF3KO cells. **G** Flow cytometry results showing the surface levels of MHC-I using an antibody conjugated to the FITC fluorophore. The mean fluorescence intensities (MFI) were compared between each condition and plotted on a bar graph. **H** Kaplan-Meier plot showing the survival of mice in four groups: NSG mice injected with ITB1 (solid pink line), NSG mice injected with TRAF3KO cells (dotted pink line), WT mice injected with ITB1 (solid cyan line), and WT mice injected with TRAF3KO cells (dotted cyan line).

We then validated the potential role of TRAF3 as a tumor-immune suppressive mediator by injecting TRAF3KO1 (henceforth referred to as TRAF3KO) and ITB1 cells into NSG and WT mice and monitoring ascites development and the survival of the mice. In NSG mice, there was no significant difference in the growth delay of tumors bearing TRAF3KO (33 days) or ITB1 (29 days) tumors, indicating that the tumors developed at similar rates in the absence of an active immune system. In contrast, in WT mice, the median survival of mice bearing ITB1 tumors was 38 days, while mice bearing TRAF3KO tumors survived for almost three times as long—a median of 98 days (Fig. 2H). These survival differences in WT but not NSG mice further support the hypothesis that TRAF3 acts an immune-suppressive modulator in ITB1 OC cells, most probably by controlling the levels of type-I IFN.

### TRAF3 suppression in OC cells induces an IgA-mediated B cell immune response

To investigate the potential mechanisms by which immune cells eliminate or reduce the growth of TRAF3KO tumors, we injected ITB1 or TRAF3KO cells i.p. into WT mice, washed out the abdominal cavity two weeks after injection, and characterized the immune populations in the washing solution by flow cytometry. Analysis of the myeloid population, determined by the expression of CD45^+^CD 11b^+^, showed a decrease in mice bearing TRAF3KO tumors compared to ITB1 tumors. Classification of the myeloid subsets stained with F4/80, CD11c, and LY6C/G showed that the percentage of macrophages (CD45^+^CD11b^+^F4/80^+^) was significantly lower in TRAF3KO tumors, while the percentages of other populations, such as dendritic cells (CD45^+^CD11b^+^CD11c^+^MHC-II^+^), monocytic cells (CD45^+^CD11b^+^Ly6C^+^Ly6G^-^), and polymorphonuclear (CD45^+^CD11b^+^Ly6C^-^Ly6G^+^) myeloid-derived suppressor cells (MDSCs), were not significantly different between the two groups of mice (Fig. 3A). In the lymphocyte population, there were no differences in the percentages of CD3^+^, CD8^+^ T cells, and NK cells. However, we did observe increases in CD19^+^ B cells and CD4^+^ T cells (Fig. 3B).

**Fig. 3.**
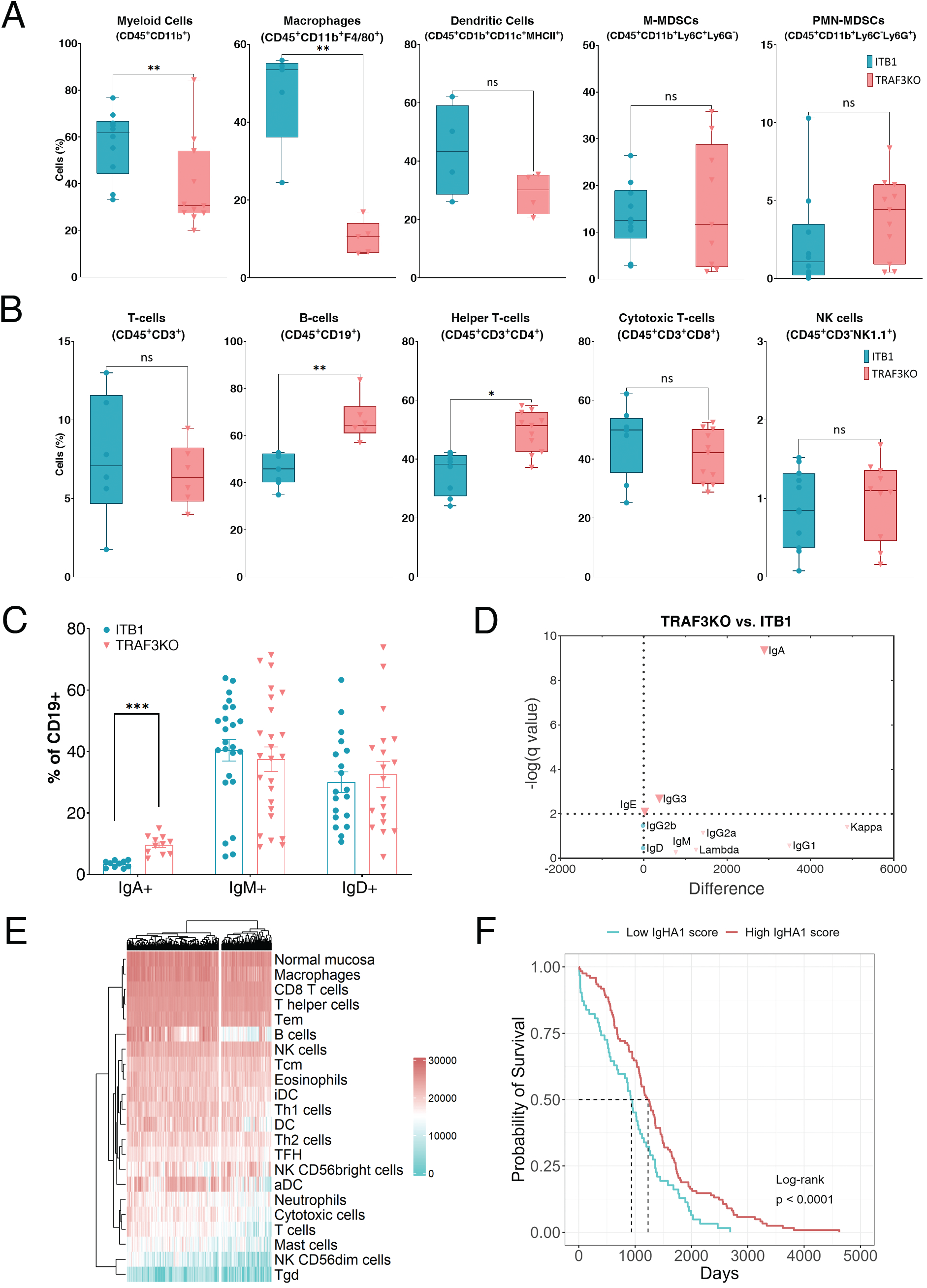
TRAF3 knockout (TRAF3KO) cells alter the immune response. **A** Profile of total myeloid cells (CD45^+^CD11b^+^), macrophages (CD45^+^CD11b^+^F4/80^+^), dendrites (CD45^+^CD11b^+^CD11c^+^MHCII^+^), monocytic (CD45^+^CD11b^+^Ly6C^+^Ly6G-), and polymorphonuclear (CD45^+^CD11b^+^Ly6C-Ly6G^+^) MDSCs in ITB1- (cyan) and TRAF3KO (pink)-injected mice. **B** Profile of total T cells (CD45^+^CD3^+^), B cells (CD45^+^CD19^+^), helper T-cells (CD45^+^CD3^+^CD4^+^), cytotoxic T-cells (CD45^+^CD3^+^CD8^+^), and NK cells (CD45^+^CD3-NK1.1^+^) in ITB1-injected (cyan) and TRAF3KO-injected (pink) mice. **C** Expression of immunoglobulin markers (IgA, IgD, IgM) on the surface of B cells in ITB1-and TRAF3KO-injected mice. **D** Volcano plot of a multiple t-test analysis comparing TRAF3KO and ITB1 washes. The plot shows the difference (x-axis) and p-value (y-axis) of the mean fluorescence intensity of each immunoglobulin scanned with an Ig isotyping array, **E** Heatmap showing that the ovarian cancer TCGA cohort is clustered according to the B cell pathway. **F** Survival curve showing the difference between patients with high (cyan) and low (pink) levels of IGHA1 in the TCGA data.

To better characterize the B cell population in mice bearing TRAF3KO tumors, we compared the maturation of B cells by co-staining CD19 with the surface forms of IgM, IgA, and IgD. In these three immunoglobulin subclasses, there was significantly higher surface expression of IgA on B cells in the ascites of mice bearing TRAF3KO tumors compared to those bearing ITB1 tumors (Fig. 3C). Notably, there were no significant differences in the expression of the costimulatory protein CD40 or the percentage of regulatory B cells determined by the intracellular expression of IL-10 (Fig. S3A). To confirm the increased expression of CD19^+^IgA^+^ cells in mice bearing TRAF3KO tumors, we used an immunoglobulin isotyping array to compare the levels of seven different immunoglobulins in the peritoneums of mice injected with ITB1 or TRAF3KO cells. Three of the seven immunoglobulins were more abundant in the peritoneal cavities of TRAF3KO-injected mice than in those of ITB1-injected mice, with IgA being the most significantly expressed immunoglobulin (Fig. 3D and Fig. S3B and C).

To explore whether our findings have any clinical relevance in OC patients, we examined the gene expression profiles and clinical data of 380 high grade serous OC cases from the Cancer Genome Atlas (TCGA) portal. These profiles were subjected to single-sample gene set enrichment analysis against an immune gene set of 28 different immune cell types [37]. In this analysis, we visualized the associations between the TCGA samples and the different data sets and split the samples by k-means clustering, creating two distinct clusters (Fig. 3E and Fig. S3D). We noticed that the factor that differed to the greatest extent between the two clusters was the B cell signature (Fig. S3E) and that one of the major genes in the B cell signature was immunoglobulin heavy constant alpha 1 (IGHA1), which is part of both the monomeric form of IgA and the secretory dimeric IgA complex [38]. Comparing the survival probability of patients based on their B cell signature score showed that the median survival of patients with a high score was significantly longer than that of patients with a low score (Fig. S3F), and much longer when differentiating the patients by expression levels of IGHA1 (Fig. 3F). These results further support the hypothesis that the enrichment of the TME with B cells and secreted IgAs may reflect an enhanced overall anti-tumor immune response, which prolongs the survival of OC patients.

### IgA-mediated B cell immune response is affected by the commensal microbiome

IgA-secreting B cells are key players in maintaining a symbiotic relationship between the host’s immune system and the commensal microbiome [39]. Disruption of this symbiotic relationship and hence of the interaction between bacteria and B cells has been implicated in several diseases, including some cancers [40,41]. To study the effect of bacteria on the antitumor B cells response, we depleted the microbiome by treating WT mice with a cocktail of four different antibiotics (ABX), as described in the Methods section. We verified the microbiome depletion by alterations in the cecum size [42,43] and by monitoring the growth of colonies isolated from the feces of treated and untreated mice on blood agar plates (Fig. S4A and B). We then injected the ABX-treated mice with TRAF3KO cells and monitored their survival in comparison with untreated mice injected with ITB1 or TRAF3KO cells. ABX treatment shortened the median survival of TRAF3KO-injected mice to 35 days, compared to 92 days for untreated TRAF3KO-injected mice (Fig. 4A). To determine the response of the immune system to ABX treatment, we collected and profiled the immune cells from the abdominal cavities of ABX-treated TRAF3KO-injected mice, compared to those from the abdominal cavities of ITB1- or TRAF3KO-injected untreated mice. For TRAF3KO-injected mice, the comparison showed overexpression of total CD19^+^ cells, and particularly CD19^+^MHC-II^+^ cells, in the untreated mice but not in the ABX-treated mice (Fig. 4B). We then looked for associations between the abundance of IgA-coated bacteria and the expression of TRAF3. To this end, we isolated and purified IgA-coated bacteria from the peritoneal washes of both TRAF3KO-injected and ITB1-injected mice, as previously described [44]. We then applied 16S sequencing to characterize the taxa-specific coated bacteria of each group of mice. We found higher abundance of two genera of IgA-coated bacteria (*Flavobacterium* and *Novosphingobium*) in the ITB1 group and one genus (*Sphingobium*) in the TRAF3KO group (Fig. 4C and Fig. S4C). Finally, using the TCGA microbiome database [45], we looked for the same associations in humans regarding IGHA1 expression. We found two bacterial genera that correlated positively and two bacterial genera that correlated negatively with IGHA1 expression (Fig. 4D).

**Fig. 4.**
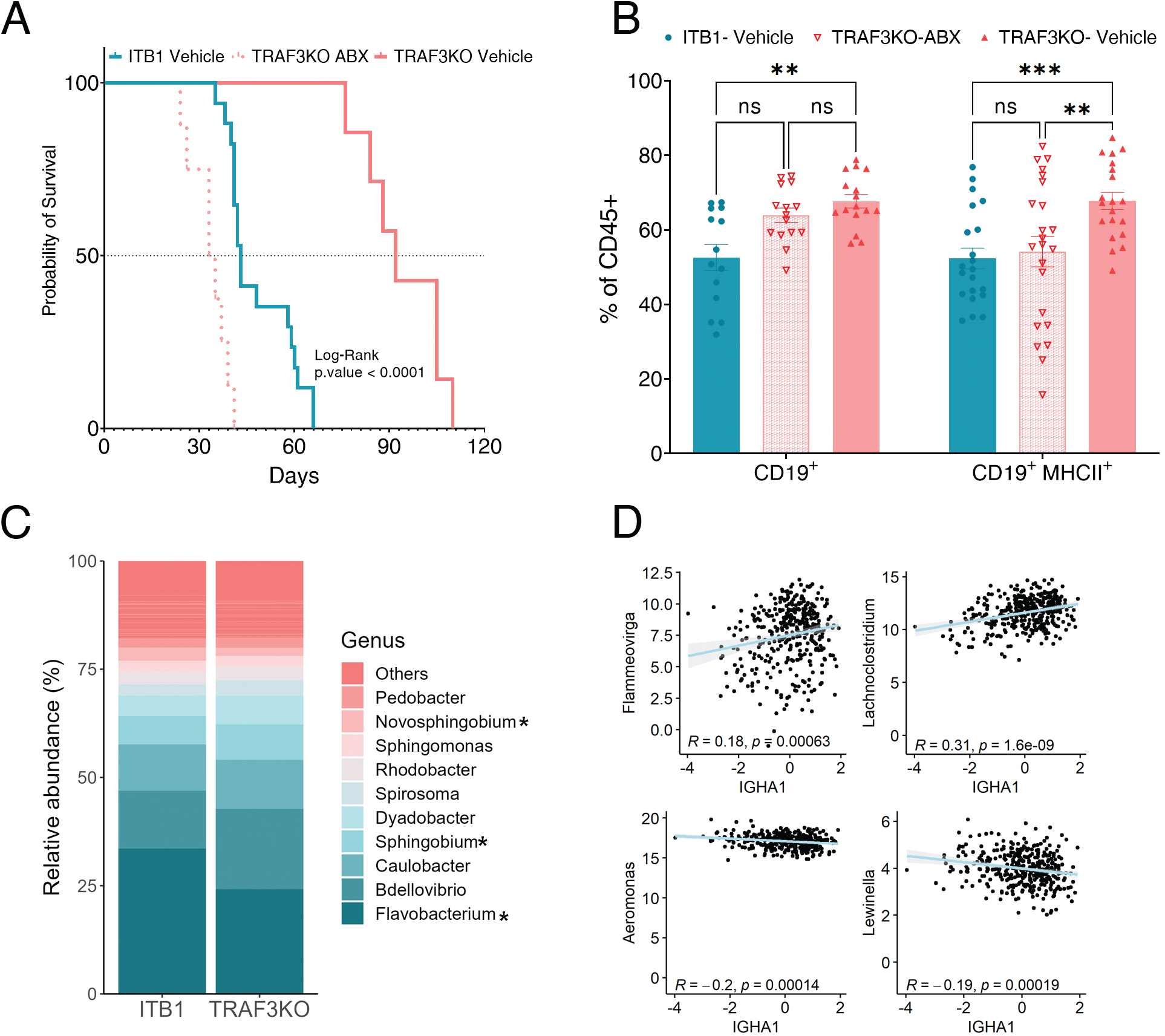
Bacteria are necessary for the B cell anti-tumor response. **A** Kaplan-Meier plot showing the survival of mice in four groups: untreated WT mice injected with ITB1 (cyan line), untreated WT mice injected with TRAF3KO cells (pink line), antibiotic (ABX)-treated WT mice injected with ITB1 (cyan dashed line), and ABX-treated WT mice injected with TRAF3KO cells (pink dashed line). **B** Bar graph representing the percentage of CD19+ cells and MHCII+ cells in ITB1 (cyan circles), TRAF3KO (pink triangles), and ABX-treated TRAF3KO- (pink open triangles) injected mice. **C** Bar plot showing the abundance of each genus in each group. **D** Correlation curves showing the positive correlation of *Flammeovirga* and *Lachnoclostridium*, and the negative correlation of *Aeromonas* and *Lewinella*, with IGHA1 expression.

Overall, these results emphasize the contribution of bacteria in the TME to an enhanced (or otherwise) anti-tumor B cell response.

### IgA-seq of ascites can distinguish ovarian cancer patients

To evaluate the association between microbial communities and IgA in human OC, we examined, as a proof of concept, 11 ascites samples from OC patients and 4 ascites samples from patients with cirrhosis. First, we isolated the total bacterial DNA from each sample, and then amplified and sequenced the 16S rRNA. After ruling out contaminations, as described in the Methods section, we analyzed the obtained sequencing data and estimated the diversity and abundance of microbial communities in each group. Using the Chao1 model, we did not observe significant differences in the α-diversity between OC and cirrhosis patients (Fig. 5A). An ordination plot based on Bray-Curtis dissimilarities (β-diversity) partially revealed distinct microbial compositions that differed between OC and cirrhosis patients (Fig. 5B). To search for differentially abundant taxa in the ascites microbiome of OC versus cirrhosis (Fig. 5C), we used the Deseq2 method to run an enrichment analysis. As ascites samples are considered low biomass samples and some amplicon sequence variants (ASVs) were found only in one sample, we ran the enrichment analysis once with ASVs expressed in more than one sample and once without a sample threshold. The enrichment analysis of the former revealed significant differences at the genus level, with one genus, *Reyranella*, being enriched in cirrhosis and one genus, *Brevundimonas*, being enriched in OC, with a log2 fold change of > 1.2 (Fig. 5D). Analysis without threshold revealed three and five significant ASVs in cirrhosis and OC, respectively (Fig. S5A).

**Fig. 5.**
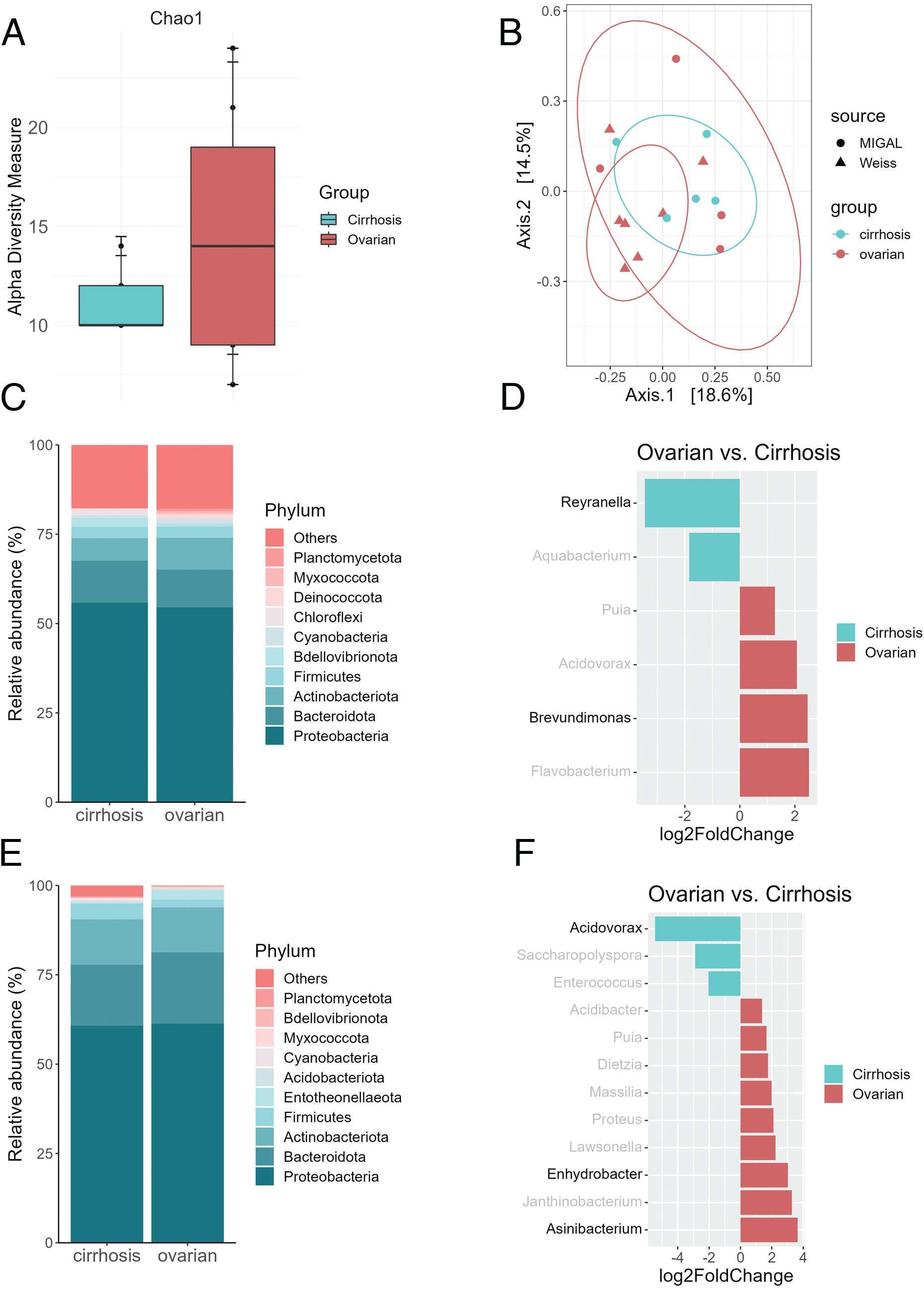
Ascites IgA-seq can differentiate ovarian cancer patients from cirrhosis patients. **A** Alpha diversity analysis using the Chao1 model shows species richness in each group. Cyan samples are from cirrhosis patients, and pink samples, from ovarian cancer (OC) patients. **B** Beta diversity analysis shows the differences in species richness between groups. **C** Bar plot showing the abundance of each phylum in the ascites of each group. **D** Bar plot showing the log2 fold change (x axis), discovered using Deseq2, of genera expressed at least in two or more patients. Genera whose expression was higher in cirrhosis or OC are shown in cyan or pink, respectively. Genera written in black are significantly expressed in one of the groups, while those written in gray are not significantly expressed. **E** Bar plot showing the abundance of each phylum in the IgA-coated bacteria of the ascites of each group. **F** Bar plot showing the log2 fold change (x axis), discovered using Deseq2, of IgA-coated genera expressed at least in two or more patients. Genera whose expression was higher in cirrhosis or OC are shown in cyan or pink, respectively. Genera written in black are significantly expressed in one of the groups, while those written in gray are not significantly expressed.

As we found an increase of IgA and infiltration of B cells in mice injected with TRAF3KO cells, we also isolated IgA-coated bacteria from the crude ascites from OC and cirrhosis patients, as described before [44], and amplified and sequenced the 16S rRNA. Analysis of the ASVs showed a complete shift in the relative abundance of taxa at the phylum level in OC compared to cirrhosis (Fig. 5E). The enrichment analysis revealed significant differences at the genus level, with one genus, *Acidovorax*, enriched in cirrhosis and two genera, *Enhydrobacter* and *Asinibacterium*, enriched in OC (Fig. 5F). Without a threshold, we found six ASVs significantly enriched in cirrhosis and five ASVs significantly enriched in OC (Fig. S5B). These results indicate that ascites contain IgA-coated bacteria and the differences in IgA-coated bacteria between cirrhosis and OC suggest potential role(s) of B cells and IgA in controlling disease progression.

## Discussion

In this study, we identified TRAF3 as a potential immunomodulator in OC cells. The involvement of TRAF3 in cancer pathogenesis has been extensively studied (reviewed in [46]) and, while most evidence identifies TRAF3 as a tumor suppressor, as found in an HPV-positive head and neck cancer cohort [47], TRAF3 was recently found to have an immune-suppressive role and to limit the response to immune checkpoint blockade in melanoma [35]. In contrast to previous findings in solid cancers that TRAF3 deficiency leads mainly to the activation of both canonical and noncanonical NF-κB pathways [35,47], we showed that knocking out TRAF3 in the ITB1 cell line did not affect NF-kB pathway activation, but rather hyperactivated the type-I IFN pathway. These contrasting findings emphasize the notion that TRAF3 functions differently in different cell types, particularly when it mediates the activity of type-I IFN [48]. Furthermore, the link between type-I IFN hyperactivation and TRAF3 can be dictated by other factors, such as the inactivation of BRCA1. DNA repair defects due to BRCA1/2 mutation induce immune signaling through the cGAS/STING/type-I IFN pathway [49]. TRAF3 inactivation may enhance the activation of STING and type-I IFN pathways in BRCA1-mutated models, but may not be sufficient for it by itself in BRCA1-wild-type models.

Type I IFNs have been shown to directly affect the proliferation, apoptosis, and other functions of tumor cells and also to indirectly affect these functions by increasing MHC-I expression [50]. Our study showed that TRAF3KO cells and ITB1 cells cultured with IFNs overexpressed MHC-I, consistent with the findings of other studies [35]. Higher levels of MHC-I enhance the expression of antigens, leading to increased recognition of antigens by antigen-presenting cells (APCs) and antigen presentation to cytotoxic T cells [51]. While we did not observe changes in the levels of cytotoxic CD8^+^ T cells, we did observe changes in the levels of macrophages and B cells in the mice bearing TRAF3KO tumors. These mice showed decreased levels of macrophages, which have previously been linked to the promotion of tumor metastasis, to enhanced angiogenesis and chemoresistance, and to poor prognosis in OC patients [52]. Likewise, the presence of plasma and memory B cells, which are often associated with improved prognosis in OC patients [11,53,54], were increased in TRAF3KO tumor-bearing mice compared to mice bearing ITB1 tumors.

The anti-tumor B cell response is largely achieved through the production of tumorspecific immunoglobulins that target tumor antigens and facilitate the recruitment and priming of APCs, leading to immune cytotoxic responses [11,55]. In our model, we observed excess production and secretion of IgA in TRAF3KO tumor-bearing mice, further emphasizing the anti-tumor role of B cells. B cell activation and differentiation can be triggered through either T-dependent or T-independent mechanisms. In both cases, the initial signal that initiates the activation cascade relies on the recognition of antigens via the B cell receptor in T-dependent activation, or via Toll-like receptors in T-independent activation [56]. These antigens are typically derived from foreign proteins, polysaccharides, or metabolites from viral, fungal, or bacterial sources [57]. Therefore, in our study, we sought to investigate the correlation between the expression and function of IgA, bacterial genera, and survival in patients with OC. We found that the presence of certain bacterial genera correlated with the expression of IGHA1 in OC patients. We demonstrated experimentally in mice that antibiotics could prevent B cell-mediated growth delay of TRAF3KO OC cells.

The presence of microbiota in the TME has been shown to play a role in many malignancies, either directly or indirectly through secreted metabolites. While this link has been studied primarily in the context of colorectal cancer using stool samples (as reviewed in [58]), some studies have linked several bacterial genera to OC in humans [15,16,59] and mice [60]. Although our cohort of patients was small, we found three genera in significant abundance in OC ascites –*Brevundimonas*, *Enhydrobacter*, and *Asinibacterium*. These genera have previously been detected in ascites fluid and even in peritonitis [61–63], making their presence in the ascites of OC patients plausible.

We also characterized the level of IgA coating on these bacteria, since bacteria–IgA interactions are known to have a significant impact on the anti-tumor immune response in several ways. First, microbiota-derived MHC-I-restricted peptides can stimulate an immune response, which may cross-react with cancer antigens [64]. Second, microbiome-derived short-chain fatty acids can modulate the expression of key factors that influence B cell differentiation and antibody responses [65]. These specific immunoglobulin responses against commensal bacteria may indirectly affect tumor growth by altering the serum metabolome and cytokine repertoire [66,67] or directly engage myeloid cells through FcR-dependent antibody-dependent cell-mediated cytotoxicity (ADCC) of tumor cells [11]. Therefore, to gain a better understanding of how the secreted IgAs and changes in the commensal microbiome may affect tumor cells directly or indirectly, it may be useful to examine the levels of different FcRs and cytokines.

## Conclusions

Overall, our research has demonstrated that TRAF3 in OC cells is an immunosuppressive modulator that down-regulates MHC class I and IFN-I signaling, limits B-cell activation, and reduces the anti-tumor immunity of B cells. We further show that the presence of an intact commensal microbiome is necessary for an effective anti-tumor B cell response.

## Abbreviations

APCs: antigen-presenting cells
ASV: amplicon sequence variant
IFN: interferon
ITB1: ID8-TRP53^-/-^-BRCA1^-/-^
OC: ovarian cancer
TME: tumor microenvironment
TRAF3: tumor necrosis factor receptor-associated factor 3
TRAF3KO: ID8-TRP53^-/-^-BRCA1^-/-^-TRAF3^-/-^
WT: wild type

## Ethics approval and consent to participate

Animal experiments were conducted in compliance with protocols established by the Institutional Animal Care and Use Committee (IACUC) of Ben-Gurion University of the Negev for ensuring animal welfare and minimizing discomfort. The animal ethical clearance protocol number used for the study was IL-23-05-2020(E).

The acquisition and analysis of human ascites samples was approved by the Helsinki committee of the Rabin Medical Center and Ziv Medical Center under protocol 0450-16-RMC and 201913481, respectively. Informed consent was obtained from all participants prior to their recruitment to the study.

## Consent for publication

Not applicable.

## Availability of data and materials

Datasets generated and analyzed during the current study are available from the corresponding author upon reasonable request.

## Authors’ contributions

JZ, YM, RDM, YKC, ZS, JM, JMG and ME were involved in experiment design and execution. JZ, ME, YM, and JGM wrote the manuscript. All authors reviewed the final version.

## Acknowledgements

We would like to thank Prof. Iain A. McNeish and his members of his lab for supplying us with the ITB1 cells. We appreciate the effort of the physicians in both hospitals, Ziv and Rabin, for collecting the ascites samples of the patients.

---

**Fig. S1** Establishing a library-expressing cells. **A** WT mouse harboring ITB1-related ascites and small tumors, indicated by the white arrows or the black arrows in the IHC-stained tissues. MLN-mesenchymal lymph node. **B** Cas9-infected ITB monoclonal isolated colonies express an active (>95%) Cas9. In all flow cytometry experiments, we used a PD-1 guide and determined its expression before and after stimulation with IFN-γ. **C** Bar graph showing the number of reads per guide. Guides with the same number of reads are grouped together and indicated by the same color.

**Fig. S2** Knocking out TRAF3 does not affect the NF-κB pathway. **A** Western blot of ITB1 and TRAF3KO lysates comparing the levels of proteins of the NF-κB pathway. The levels of p65 and RelB were monitored in the nucleus, and p100 and p52 were monitored in the whole lysates. Actin and histone H3 were used as loading controls. **B** Quantitative PCR shows the mRNA levels of genes related to the NF-κB pathway (RelB, IL-6, and IκBα). Cyan bars represent ITB1, solid pink bars represent TRAF3KO1, and dotted pink bars represent TRAF3KO2. **C** MFI of the blot on the main figure, calculated using ImageJ **D** Bar graph showing the differences in mean fluorescence intensity of the Cy3 channel in the nucleus between ITB1 (cyan) and TRAF3KO cells (pink).

**Fig. S3** B cells in TRAF3KO-injected mice express and secrete more IgAs than in ITB1-injected mice. **A** Expression of the costimulatory protein CD40 and interleukin-10 (IL-10) on B cells in ITB1-injected and TRAF3KO-injected mice. **B, C** Ig isotyping array (B) and its quantification (C). **D** Scatter plot showing the differentiation between the two clusters of the heatmap. **E** Heatmap showing the average score of each cluster for each immune pathway. **F** Survival curve showing the difference between patients with high B cell score (cyan) and low B cell score (dark pink).

**Fig. S4** Effect of ABX treatment. **A** Mouse treated with ABX for 10 days and its enlarged cecum. **B** Fecal samples were collected from both ABX-treated and naïve mice and seeded on a blood agar plate. Growth of colonies can be seen only on the left side of the plate where the naïve cells was seeded. **C** Difference in the abundance of three genera, compared using the Wilcoxon signed-rank test.

**Fig. S5** Unique bacteria in the ascites of ovarian cancer patients. **A** Bar plot showing the log2 fold change (x axis), discovered using Deseq2, of genera expressed at least in one or more patients. Genera expressed more in cirrhosis or OC are in cyan or pink, respectively. Genus with black name are expressed significantly in one of the groups, while the grey ones are not significant. **B** Bar plot showing the log2 fold change (x axis), discovered using Deseq2, of IgA-coated genera expressed at least in one or more patients. Genera whose expression was higher in cirrhosis or OC are shown in cyan or pink, respectively. Genera written in black are significantly expressed in one of the groups, while those written in gray are not significantly expressed.

