## Supplemental Figures for "TRAF3 Suppression Encourages B Cell Recruitment and Prolongs Survival of Microbiome-Intact Mice with Ovarian Cancer"

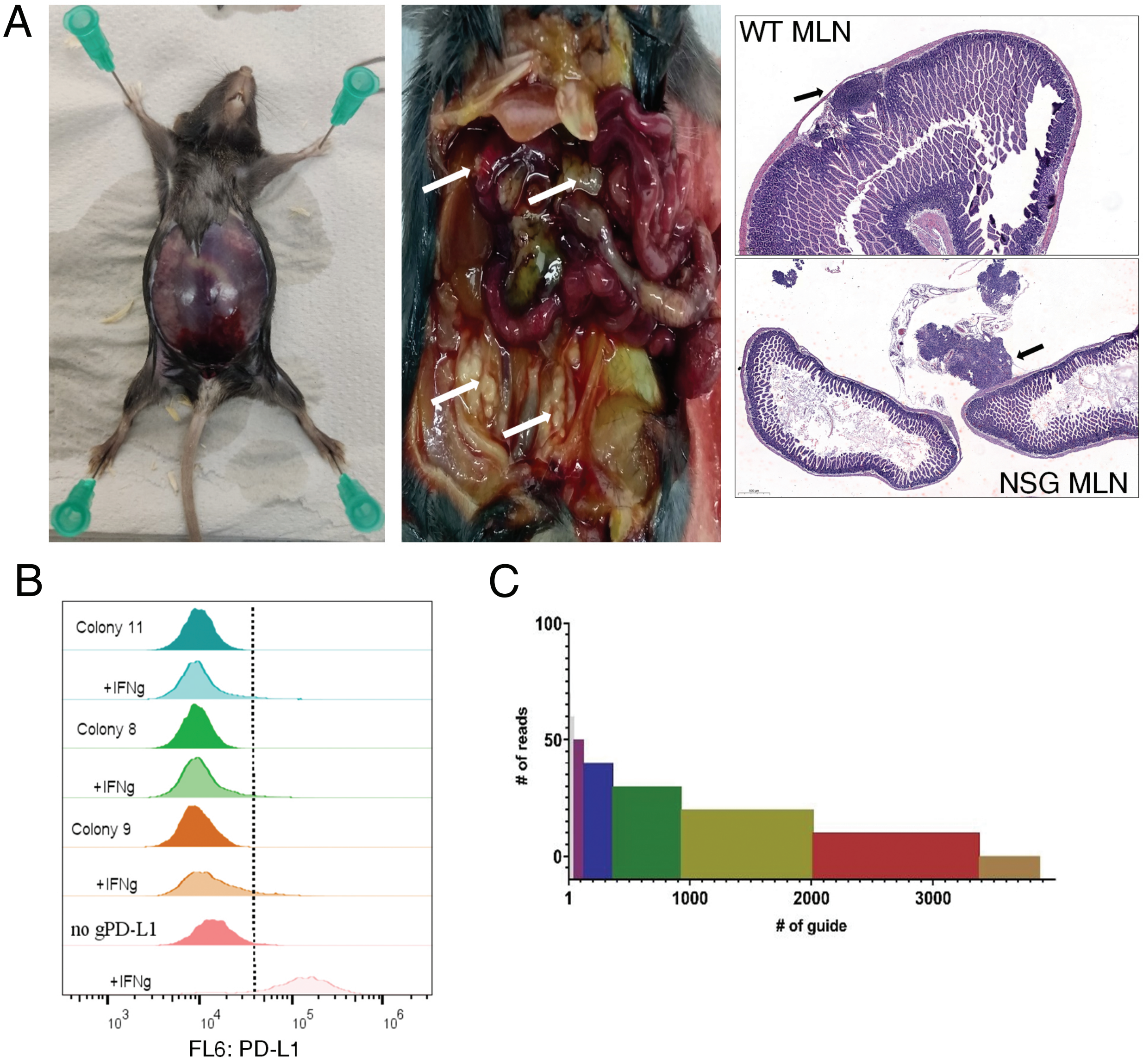
A
WT MLN
NSG MLN
B
Colony 11
+IFNg
Colony 8
+IFNg
Colony 9
+IFNg
no gPD-L1
+IFNg
10^3
10^4
10^5
10^6
FL6: PD-L1
C
100
50
0
### of reads
1
1000
2000
3000
### of guide

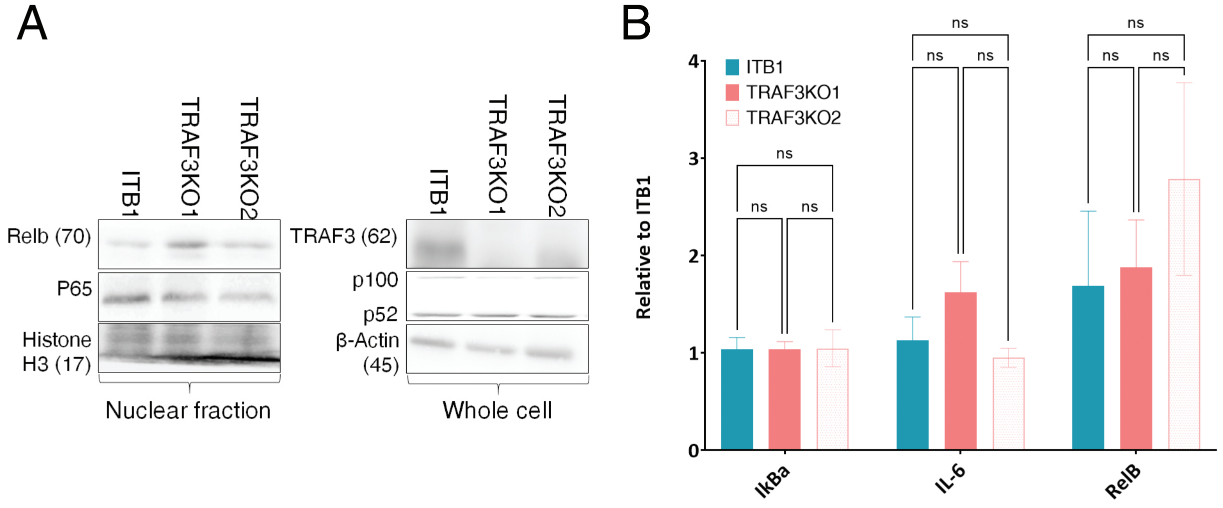


C

| | | ITB1 | TRAF3KO1 | TRAF3KO2 |
|---|---|---|---|---|
| TRAF3 | Total | 313.6 | 0.0 | 0.0 |
| | Cytoplasmic | 714.7 | 0.0 | 0.0 |
| | Nuclear | 267.1 | 0.0 | 0.0 |
| pTBK1 | Total | 4592.4 | 2011.5 | 2452.0 |
| | Cytoplasmic | 1225.4 | 2068.0 | 2081.5 |
| | Nuclear | 1367.8 | 1015.7 | 1097.1 |
| TBK1 | Total | 1124.5 | 830.2 | 937.7 |
| | Cytoplasmic | 2993.7 | 4696.1 | 5547.5 |
| | Nuclear | 0.0 | 0.0 | 0.0 |
| STING | Total | 4334.2 | 6243.8 | 4812.9 |
| | Cytoplasmic | 910.6 | 1533.0 | 1822.5 |
| | Nuclear | 1309.7 | 1774.2 | 2377.6 |
| β-Actin | Total | 3718.6 | 3290.4 | 2661.7 |
| | Cytoplasmic | 3159.8 | 2467.2 | 2737.9 |
| | Nuclear | 2479.8 | 2064.1 | 2405.9 |
| Histone H3 | Total | 816.4 | 608.4 | 562.9 |
| | Cytoplasmic | 1158.9 | 1107.5 | 438.5 |
| | Nuclear | 21651.0 | 22899.9 | 21956.8 |

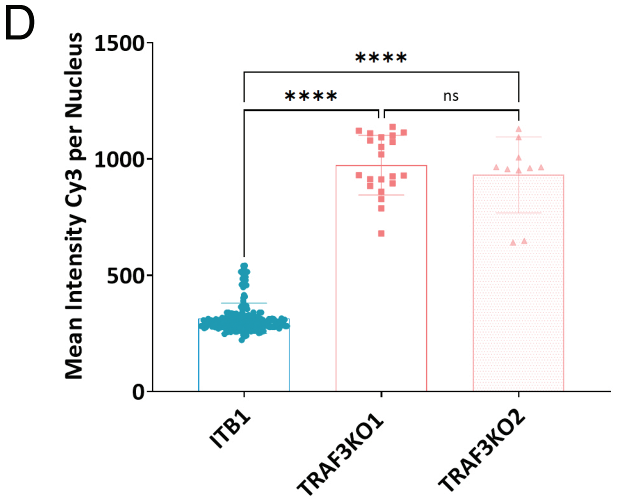

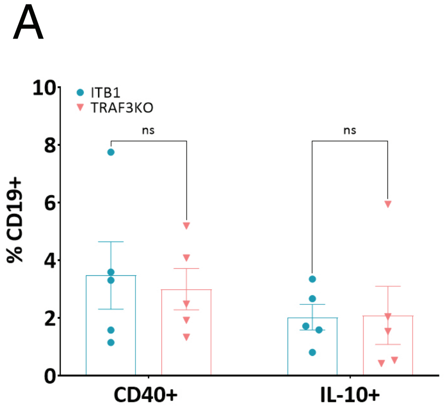


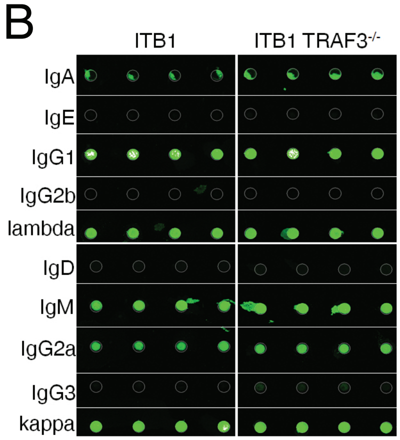


C

| | P value | Mean of ITB1 | Mean of ITB1 TRAF3-/- | Adjusted P Value |
|---|---|---|---|---|
| IgA | 0.01> | 2769 | 5157 | 0.01> |
| IgD | 0.34 | 192.8 | 176.2 | 0.71 |
| IgE | 0.01> | 60.38 | 87.75 | 0.02 |
| IgM | 0.68 | 16933 | 17703 | 0.71 |
| IgG1 | 0.23 | 33466 | 36961 | 0.65 |
| IgG2a | 0.05 | 6398 | 7821 | 0.24 |
| IgG2b | 0.02 | 66.75 | 48.19 | 0.11 |
| IgG3 | 0.01> | 252.4 | 630.0 | 0.01> |
| Lambda | 0.46 | 16645 | 17899 | 0.71 |
| Kappa | 0.02 | 20318 | 25200 | 0.14 |

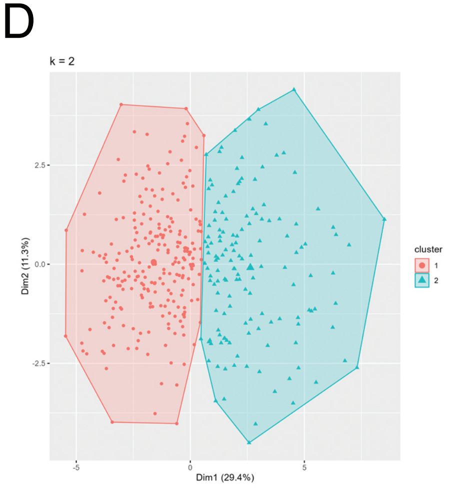


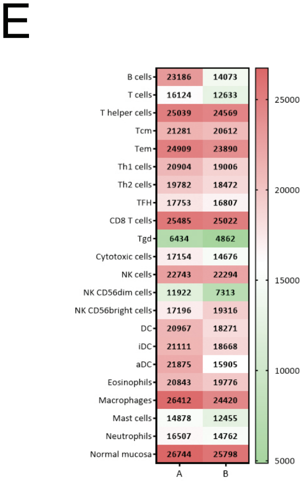


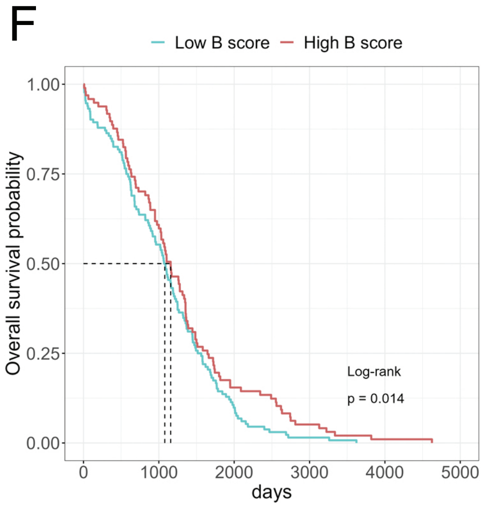

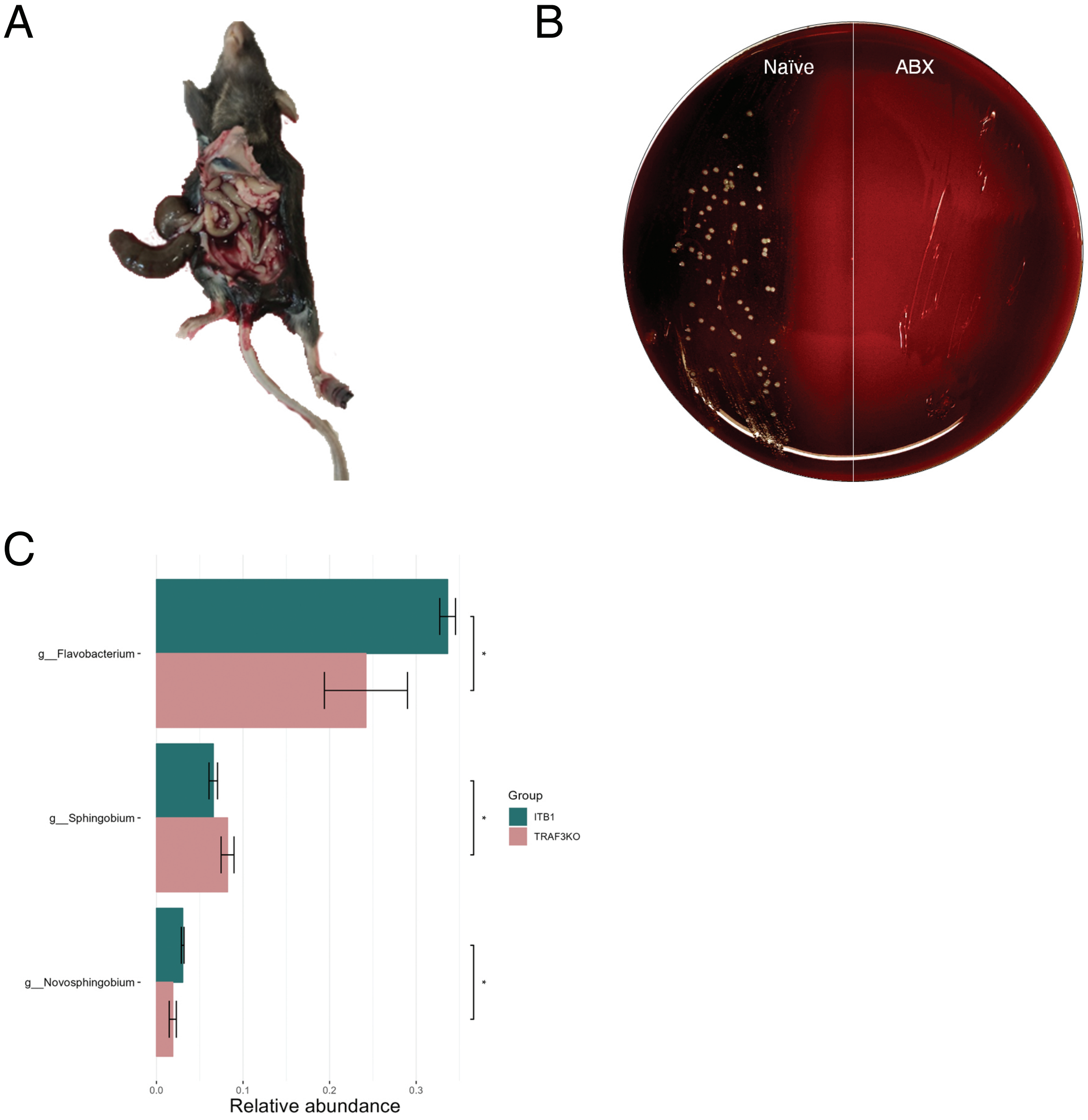
A
B
Naïve
ABX
C
g__Flavobacterium
g__Sphingobium
g__Novosphingobium
*
*
*
Group
ITB1
TRAF3KO
0.0
0.1
0.2
0.3
Relative abundance

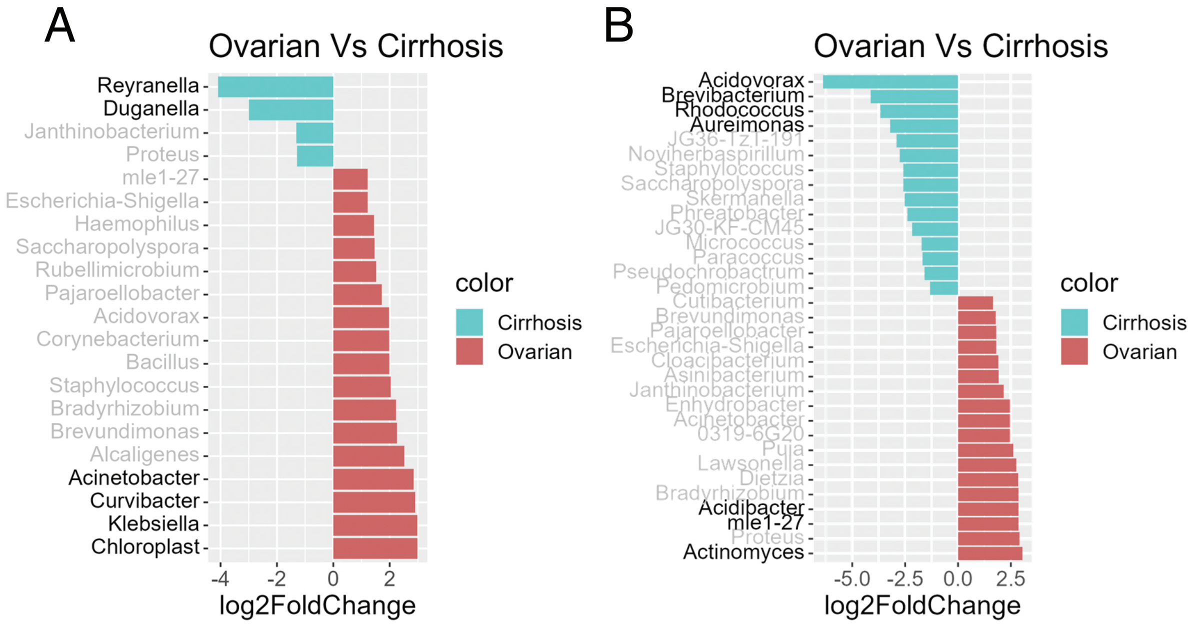
A
Ovarian Vs Cirrhosis
Reyranella
Duganella
Janthinobacterium
Proteus
mle1-27
Escherichia-Shigella
Haemophilus
Saccharopolyspora
Rubellimicrobium
Pajaroellobacter
Acidovorax
Corynebacterium
Bacillus
Staphylococcus
Bradyrhizobium
Brevundimonas
Alcaligenes
Acinetobacter
Curvibacter
Klebsiella
Chloroplast
-4
-2
0
2
log2FoldChange
color
Cirrhosis
Ovarian
B
Ovarian Vs Cirrhosis
Acidovorax
Brevibacterium
Rhodococcus
Aureimonas
JG36-TzT-191
Noviherbaspirillum
Staphylococcus
Saccharopolyspora
Skermanella
Phreatobacter
JG30-KF-CM45
Micrococcus
Paracoccus
Pseudochrobactrum
Pedomicrobium
Cutibacterium
Brevundimonas
Pajaroellobacter
Escherichia-Shigella
Cloacibacterium
Asinibacterium
Janthinobacterium
Enhydrobacter
Acinetobacter
0319-6G20
Puia
Lawsonella
Dietzia
Bradyrhizobium
Acidibacter
mle1-27
Proteus
Actinomyces
-5.0
-2.5
0.0
2.5
log2FoldChange
color
Cirrhosis
Ovarian
